# Integrating explainable AI with multiomics systems biology and EHR data mining for personalized drug repurposing in Alzheimer’s disease

**DOI:** 10.1101/2025.03.24.644676

**Authors:** Mohammadsadeq Mottaqi, Pengyue Zhang, Lei Xie

## Abstract

Alzheimer’s disease (AD) is characterized by region- and patient-specific molecular heterogeneity, which hinders therapeutic design. In this study, we introduce PRISM-ML (PRecision-medicine using Interpretable Systems and Multiomics with Machine Learning), an open-source integrated analysis pipeline that combines interpretable machine learning with systems biology and electronic health record (EHR) data mining to elucidate the molecular diversity of AD and predict promising drug repurposing opportunities. First, we integrated and harmonized transcriptomic (bulk RNA-seq) and genomic (genome-wide association study) data from 2105 brain samples, each with matched data from the same individual (1363 AD patients, 742 controls; nine tissues), sourced from three independent studies. Random forest classifiers with SHapley Additive exPlanations (SHAP) identified patient-specific biomarkers; unsupervised clustering resolved 36 molecularly distinct “subtissues” (clusters of samples); and gene-gene co-expression networks prioritized 262 high-centrality bottleneck genes as putative regulators of dysregulated pathways. Next, knowledge graph-based drug repurposing predicted six FDA-approved drugs that simultaneously target multiple bottleneck genes and multiple AD-relevant pathways. Notably, in a large U.S. de-identified insurance-claims database (n = 364733), exposure to promethazine, one of the candidate drugs, was associated with a 57–62 % lower incidence of AD versus an active antihistamine comparator (adjusted hazard ratio 0.38; inverse-probability weighted 0.43; both p < 0.001), providing real-world support for its repurposing potential. In summary, PRISM-ML, as an explainable multi-omics analysis pipeline, is readily transferable to other complex diseases, advancing precision medicine.

## Introduction

Alzheimer’s disease (AD) is a progressive neurodegenerative disorder that affects millions of people worldwide, yet its underlying molecular mechanisms remain elusive (*1*). Current drugs provide only symptomatic relief and fail to target the disease itself, largely because of the profound molecular heterogeneity of AD across patients and brain tissues (*2*). The advent of multiomics technologies, which integrate data from real patients—such as transcriptomics, genomics and proteomics—offers a powerful means to unravel this complexity. Integrating multiple omics layers is critical for capturing the full spectrum of molecular dysregulation in AD, revealing interactions missed by single-omics studies. However, translating these rich datasets into actionable therapeutic strategies remains a significant challenge, underscoring the need for innovative approaches that go beyond traditional methods (*1*).

Previous research has often relied on single-omics studies, which, while informative, capture only a fraction of the molecular landscape of AD (*3*). More recent efforts have applied machine learning (ML) and deep learning (DL) to multiomics data, identifying biomarkers, gene sets, or potential drug targets (4). However, a major drawback of these ML and DL models is their "black box" nature, which makes it difficult to understand how predictions are made or to gain insights into the underlying disease mechanisms (*5*).

A notable exception is a recent study (*6*) that combined weighted gene co-expression network analysis (WGCNA) (7) for feature selection with SHapley Additive exPlanations (SHAP) (*8*) to improve model interpretability. By using SHAP to measure feature importance, this study made the ML decision-making process more transparent and developed a global 10-gene diagnostic panel for AD on the basis of a single GEO microarray cohort. This work highlights the promise of interpretable ML in advancing AD research. Building on this, our analysis pipeline, PRISM-ML (PRecision-medicine using Interpretable Systems and Multiomics with Machine Learning), employs SHAP (*8*) to deliver explanations for individual patients, revealing both the magnitude and direction of contribution of each gene to AD predictions (classifications). This directional insight, which is absent in traditional feature importance methods such as the mean-decrease-in-impurity of random forest (RF), enhances interpretability and supports the identification of patient-specific biomarkers.

Moreover, many studies propose drugs for AD treatment without robust validation, often relying on cell lines or animal models that may not reflect human disease (9). Additionally, patient-specific details—crucial for personalized medicine—are rarely considered (9), leaving a gap between computational predictions and clinical relevance.

To overcome these limitations and bridge the gap in clinical application, we developed PRISM-ML, a novel open-source analysis pipeline that integrates interpretable ML, statistical analyses, network biology, and electronic health record (EHR) data mining aiming to transform large-scale multiomics data into actionable insights into the molecular complexity of AD and patient responses in the real world.

Leveraging matched transcriptomic (bulk RNA-seq) and genomic (genome-wide association study (GWAS)) data from 2105 postmortem brain samples (1,363 AD, 742 controls) across nine brain tissues, PRISM-ML provides a holistic view of the dysregulated disease pathways. We clustered these samples into 36 distinct "subtissues" (clusters) on the basis of gene expression profiles, identifying patient-specific and subtissue-specific biomarkers and genetic drivers. Subtissue-specific gene‒gene interaction networks prioritized 262 "bottleneck" genes—key regulators of dysregulated pathways—as drug targets. Leveraging a biomedical knowledge-graph approach, we predicted six FDA-approved, multitarget drugs capable of modulating these pathways.

While reliant on postmortem samples, our approach is validated through a pharmacoepidemiologic study using healthcare data from 364733 individuals to determine its clinical relevance. These findings suggest new therapeutic strategies through the identification of patient-specific biomarkers, subtissue-level heterogeneity, and multitarget drug repurposing which potentially paves the way for personalized treatments tailored to individual molecular profiles. Beyond AD, PRISM-ML offers a scalable analysis pipeline adaptable to other complex diseases.

This manuscript is organized as follows: The Results section introduces the PRISM-ML analysis pipeline and presents our findings, including data processing, subtissue stratification, identification of biomarkers and bottleneck genes, drug repurposing, and pharmacoepidemiologic validation of the protective effects of promethazine. The Discussion and Conclusion sections explore the research problems, limitations, significance, and implications of our work for AD treatment. Finally, the Methods section details the technical aspects of our approach.

## Results

### 1) Overview of the PRISM-ML Pipeline

In this study, we developed PRISM-ML, a four-stage integrated analysis pipeline that fuses transcriptomic and genomic information to dissect AD heterogeneity (Figure 1). Leveraging matched bulk RNA-seq and GWAS data from 2105 postmortem samples spanning nine brain tissues (see Table 1), PRISM-ML first identifies sample-specific biomarker signatures and captures AD region-dependent molecular variation. The analysis pipeline traces dysregulated genes and pathways through network analysis and predicts repurposable drugs followed by electronic health record (EHR) data mining to validate drug effects.

**Figure 1.**
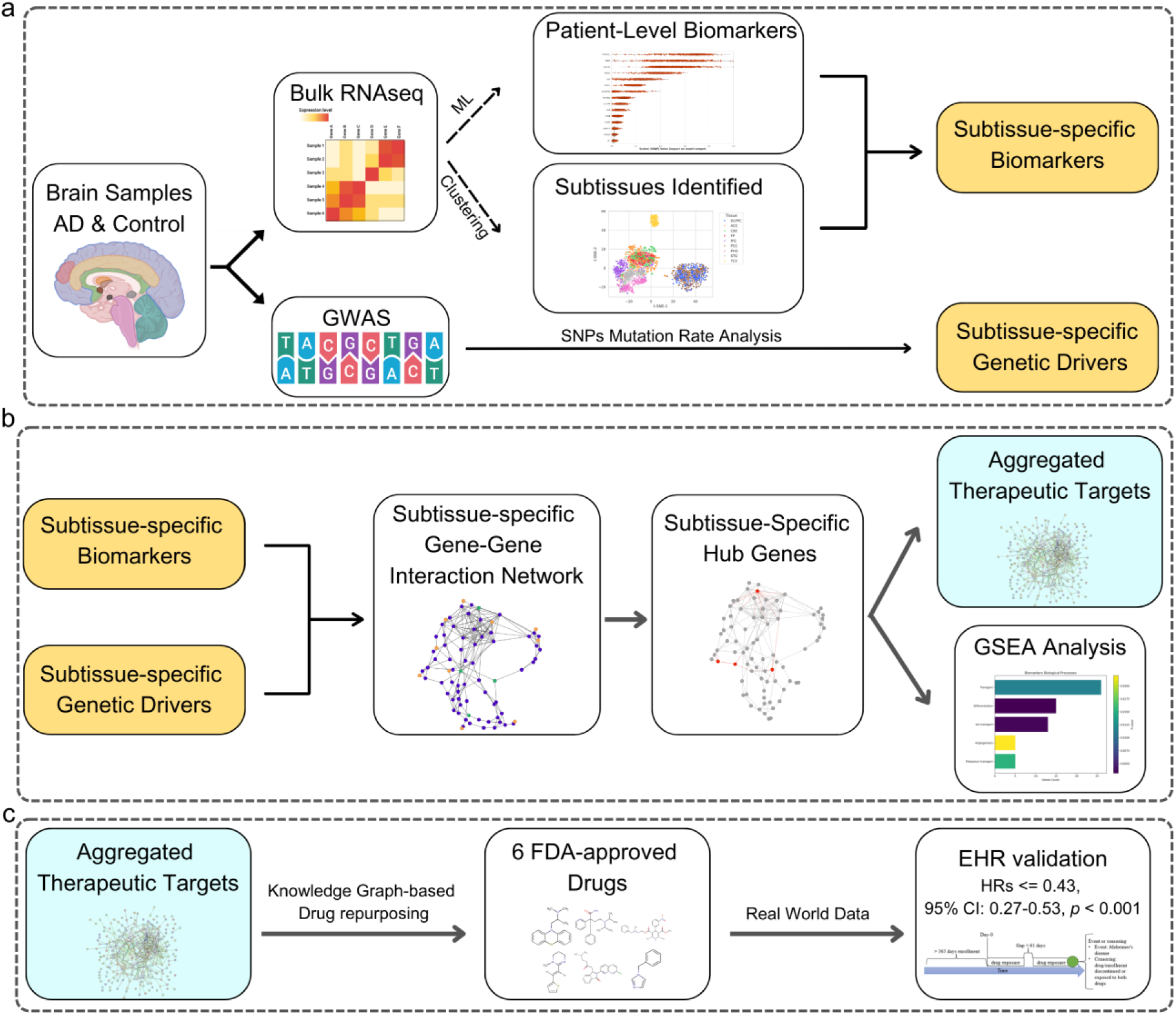
PRISM-ML: integrating interpretable machine learning, multiomics systems biology, and electronic health record (EHR) data mining for Alzheimer’s disease (AD) drug repurposing. a) Multiomics integration and subtissue identification: Bulk RNA-seq and GWAS data from 2105 postmortem brain samples (1,363 AD patients and 742 controls) across nine tissues were harmonized. A random forest classifier with SHapley Additive exPlanations (SHAP) analysis identified ≈175 patient-specific unique biomarkers per sample on average. ‘Patient-specific’ here denotes biomarkers extracted from a model trained on all other donors and interpreted for that single donor (see *Methods Section 2)*. Unsupervised clustering stratified each tissue into four molecularly distinct subtissues (36 in total). Subtissue-specific biomarkers—high-impact genes shared across samples within each subtissue cluster—were derived by intersecting patient-level biomarker sets. Mutation rate analysis of AD-associated variants revealed subtissue-specific genetic drivers. b) Subtissue-specific networks and bottleneck genes: Subtissue-specific gene‒gene interaction networks connect biomarkers and genetic drivers via critical intermediate "message-passing" genes (shortest paths). Topological metrics (e.g., betweenness centrality) prioritized 262 high-centrality bottleneck genes enriched in synaptic transmission, ion transport, and extracellular matrix organization. c) Drug repurposing and pharmacoepidemiologic study: Knowledge-graph screening flags six FDA-approved drugs (e.g., promethazine and disopyramide) that target multiple bottleneck genes. In a cohort of 364733 individuals, promethazine use was associated with 57-62% reduced AD risk (adjusted hazard ratio 0.38; inverse-probability weighted 0.43; both p < 0.001).

**Table 1.** Overall Cohort Description: Demographics, Brain Tissue Distribution, and Sample Characteristics. This table summarizes the distribution of samples and unique individuals across multiple brain tissues and studies, including the number of Alzheimer’s disease samples vs. control samples, the breakdown by sex, and total individuals per cohort.

| Tissue Name | Cohort | Total Samples | AD Samples | Control Samples | Male Samples | Female Samples | Total Subjects |
| --- | --- | --- | --- | --- | --- | --- | --- |
| ACC | ROSMAP | 265 | 171 | 94 | 90 | 175 | 265 |
| DLFPC | ROSMAP | 436 | 293 | 143 | 143 | 293 | 429 |
| PCC | ROSMAP | 250 | 151 | 99 | 97 | 153 | 249 |
| FP | MSBB | 215 | 146 | 69 | 83 | 132 | 197 |
| IFG | MSBB | 201 | 132 | 69 | 86 | 115 | 181 |
| PHG | MSBB | 226 | 155 | 71 | 84 | 142 | 168 |
| STG | MSBB | 220 | 156 | 64 | 93 | 127 | 181 |
| CBE | MAYO | 144 | 79 | 65 | 69 | 75 | 144 |
| TCX | MAYO | 148 | 80 | 68 | 66 | 82 | 148 |
| ALL TISSUES | MERGED | 2105 | 1363 | 742 | 811 | 1294 | 1962 |

Conceptually, PRISM-ML followed four major steps. First, for each sample we trained a dedicated RF classifier on the remaining 2104 samples (leave-one-out cross-validation (LOOCV) strategy). SHAP values (*8*) corresponding to features (genes) were then computed to quantify the contribution of each gene to that sample’s AD-versus-control model prediction. Genes whose SHAP value exceeded a threshold (defined in *Methods section 2*) were labeled high-impact genes (biomarkers), yielding a personalized gene set (≈175 genes on average) for each individual sample.

Second, within each brain tissue, unsupervised clustering of the AD expression profiles partitioned the samples into distinct “subtissues,” aiming to capture local molecular patterns. Consequently, a unique list of biomarkers per subtissue was identified by selecting genes that consistently showed high importance (on the magnitude of SHAP values) across all AD samples within that subtissue. In parallel, subtissue-specific genetic drivers were identified by comparing the mutation rate of well-characterized AD genes in the samples within the subtissues compared to the background population.

Third, gene-gene interaction (coexpression) networks were constructed for each subtissue. In each network, subtissue-specific biomarkers and genetic drivers were linked to each other via the shortest paths. In each filtered network, “bottleneck” (hub) genes—those with high betweenness centrality were prioritized as drug targets. These bottleneck genes are potential key mediators in dysregulated AD-related pathways because of their central roles in the network. The union of the identified bottleneck gene sets (one per subtissue) was considered the ultimate drug targets.

Finally, knowledge-graph embedding of 30 M biomedical relations between biological entities (10) predicted six FDA-approved drugs——including promethazine, disopyramide and nicardipine that potentially target multiple bottleneck genes. These repurposable drugs are promising candidates for influencing several AD-relevant pathways simultaneously, suggesting a multitarget therapeutic approach. Notably, in an independent de-identified insurance-claims cohort (n = 364 733) analyzed with an active-comparator design, exposure to promethazine was associated with a 57–62 % lower incidence of AD than the matched antihistamine cyproheptadine (adjusted hazard ratio 0.38; inverse-probability weighted 0.43; both *p* < 0.001), corroborating its therapeutic promise.

PRISM-ML advances precision medicine by unifying sample-specific/patient-specific biomarkers, subtissue-level pathology, and multitarget drug discovery into a cohesive open-source analysis pipeline. Its interpretable design addresses the "black box" limitations of conventional ML, whereas its systems biology approach suggests shared mechanisms between AD and comorbidities such as cardiovascular diseases (see *Results section 5.2*). This analysis pipeline offers a blueprint for studying complex diseases, prioritizing actionable therapeutic insights over isolated molecular findings.

### 2) PRISM-ML Identifies Patient-level AD Biomarkers

First, we compiled a multiomics dataset by merging matched bulk RNA-sequencing data and individual-level genotype data (GWAS) from the same donors in three large-scale AD brain tissue studies: the Religious Orders Study and Rush Memory and Aging Project (ROSMAP) (11), the Mount Sinai Brain Bank (MSBB) (12), and the Mayo RNA-seq study (MAYO) (13). This integration yielded 2105 postmortem brain tissue samples—1,363 classified as AD patients and 742 as controls—originating from 307 males and 528 females (see *Methods section 1*). Within each tissue, all samples originated from a single cohort; their distribution is summarized in Table 1. By ensuring that both the transcriptomic and genomic data were derived from the same individuals, this harmonized dataset provides a robust foundation for downstream analyses, capturing patient-level molecular variation across diverse brain tissues. Figure 2a presents a t-distributed stochastic neighbor embedding (t-SNE) plot visualizing the distribution of total samples in the data using 15300 genes as input features.

**Figure 2.**
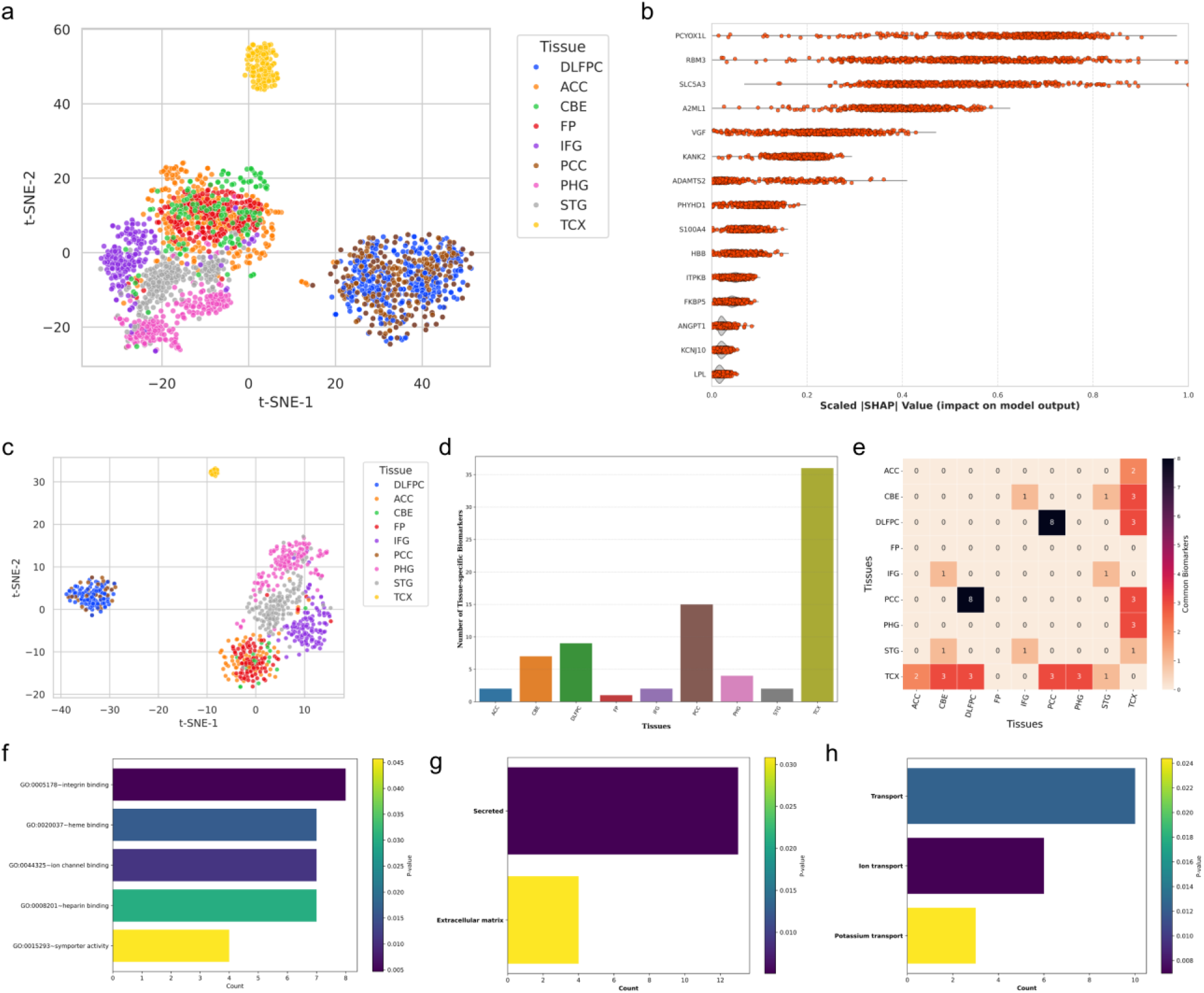
Sample-level and tissue-specific biomarkers reveal regionally conserved molecular dysregulation in Alzheimer’s disease. a) T-distributed stochastic neighbor embedding (t-SNE) visualization of all 2105 postmortem brain samples (AD patients and controls) colored by tissue of origin, illustrating the overall distribution before filtering. b) Scaled (SHapley Additive exPlanations) SHAP value distributions for representative genes, ranked by their contribution to the Random Forest models’ classification of AD vs. control. Each point corresponds to one sample, demonstrating how individual genes differentially influence the outputs of the models. c) t-SNE projection of the 720 AD samples that exceeded the average predictive score (i.e., “confidently classified” AD cases), showing the final subset used for tissue-specific biomarker analysis. d) Bar plot of the number of “common biomarkers” found in each brain tissue (i.e., high-impact genes shared by all confidently classified AD samples within that tissue). The temporal cortex (TCX) and posterior cingulate cortex (PCC) subtissues exhibit the largest sets. e) Heatmap illustrating overlaps of common biomarkers among different tissues; darker cells denote a greater degree of shared genes. f–h) Gene set enrichment analysis (GSEA) results for the union of all tissue-specific biomarkers (56 genes). (f) Enrichment in membrane-binding activities (integrin, heme, channel, heparin, symporter), (g) shows associations with secreted proteins and extracellular matrix organization, and (h) emphasizes processes involved in transport, including ion and potassium transport. Collectively, these results underscore the biological importance of the machine-learning– derived biomarkers, pinpointing pathways central to AD pathogenesis.

To identify sample-specific and ultimately patient-specific biomarkers underlying AD heterogeneity, we analyzed the gene expression profiles of the brain samples. We trained RF classifiers—using a LOOCV scheme with the top 5000 most variable genes across the dataset as features (details in *Methods section 2)*. At each iteration, a model was trained on every sample except a single test case; the withheld sample was then classified as AD or control by the model. Table 2 summarizes the strong class-specific performance of the models. Additionally, we obtained a predictive score for each sample (score ranging from 0–1) determined by the RF classifier. The score indicated the likelihood of the sample belonging to the AD group.

**Table 2.** Aggregate classification metrics from the 2105 bespoke random forest models generated in leave-one-out cross-validation. For every brain sample, a dedicated random forest classifier was trained on the remaining 2104 samples and then evaluated on the held-out sample; the SHapley Additive exPlanations (SHAP) values of the same model were used to define a sample-specific biomarker set (See *Methods section 2*) The table reports the mean F1, precision, and recall achieved by this ensemble of per-sample RF models when distinguishing AD from control tissue.

|  | F1 | Precision | Recall |
| --- | --- | --- | --- |
| AD class | 0.909 | 0.879 | 0.942 |
| Control class | 0.816 | 0.877 | 0.762 |

#### 2.1 SHAP Analysis Pinpoints Personalized Biomarker Sets

To explain each model’s prediction, we applied SHAP (8) which, for each predictive model, computes a gene-level “importance” score reflecting how much that gene/feature influences the classification decision. For every held-out sample we ranked genes by their SHAP value and flagged those genes that exceeded the cutoff (defined in *Methods section 2)* as sample-specific biomarkers (high-impact genes), indicating genes whose expression had a decisive impact on that sample’s classification.

Across the 2105 samples this procedure yielded a mean of ≈175 high-impact genes per sample, capturing unique molecular signatures that separate the diseased sample from the control (healthy) samples. Figure 2b illustrates the distribution of SHAP values for several genes across all the models, highlighting the differential impact of genes on the decisions of the models across AD samples.

To improve analytical reliability, we retained only AD samples that the model classified with high confidence—those with a predictive score > 0.82 (the mean cohort predictive score). This filter produced 720 confidently classified AD cases, excluding lower-confidence “borderline” samples without introducing systematic bias. All downstream comparisons therefore focus on this robust subset, and we refer to their high-impact gene lists as patient-specific biomarkers. Figure 2c presents a t-SNE plot visualizing the distribution of the 720 robustly classified AD cases using 5000 highly variable genes as input features. The evident tissue-level segregation in both Figure 2a and Figure 2c suggests that the RNA-seq data inherently reflect tissue-specific differences.

#### 2.2 High-SHAP Genes Convergion into Tissue-Specific Biomarker Sets

Using the 720 confidently classified AD patients, we grouped them by their tissue of origin (one of nine brain tissues). Each AD sample had its own set of high-SHAP biomarkers within each tissue. We then intersected the biomarker sets of all AD samples belonging to that tissue, identifying “common” biomarkers—genes deemed critical across all individuals within the same brain tissue. On average, we found 1 to 36 such genes per tissue, with temporal cortex (TCX) and the posterior cingulate cortex (PCC) showing the greatest overlap, reflecting regionally conserved dysregulation (Figure 2d). As shown in Figure 2e, the tissue-specific biomarker sets display varying degrees of overlap across different brain tissues, with darker cells indicating more shared genes. These findings confirm that, despite patient-level variability, there are consistently influential genes within specific brain tissues, implicating shared mechanisms in AD pathology.

#### 2.3 Tissue-specific Biomarkers Demonstrate Clear Biological Significance

To assess whether our RF–SHAP approach identified biologically meaningful genes, we combined all the tissue-specific biomarker sets—yielding a total of 56 genes across tissues—and performed GSEA via the DAVID Bioinformatics database (14). The enriched pathways and gene ontology (GO) terms included membrane-binding functions (e.g., integrin, heme, channel, symporter binding) and extracellular matrix organization (Figure 2f, g), which are pathways critical for synaptic integrity and are disrupted in AD (15). Notably, dysregulated ion transport (Figure 2h) aligns with Aβ-induced neuronal hyperexcitability (16), confirming that our biomarkers are linked to AD pathophysiology. These findings suggest that our ML analysis pipeline successfully discerns relevant molecular features rather than random signals.

### 3) Subtissue-Specific Biomarkers and Genetic Drivers Reveal Regional Heterogeneity in AD

#### 3.1 Four Subtissues per Brain Tissue Capture Finer Molecular Diversity

After identifying patient-specific biomarkers, we aimed to examine the molecular heterogeneity within each brain tissue at a more detailed level. For each of the nine brain tissues, we compiled a set of genes by combining the patient-specific biomarker sets from all samples within that tissue, taking their union to ensure that all relevant genes were included. We then applied K-means clustering to the transcriptomic profiles of these samples, using the expression levels of the compiled gene set as input features.

Using clustering evaluation metrics (detailed in *Methods section 3.1*), we determined that four stable clusters per tissue provided an optimal representation of molecular diversity. These clusters, which we term "subtissues," consist of samples with similar expression patterns. This process resulted in a total of 36 subtissues across the nine brain tissues.

#### 3.2 Subtissue-Specific Biomarker Sets Expose Localized AD Mechanisms

To identify subtissue-specific biomarkers in AD, we started with patient-specific high-impact gene sets—approximately 175 genes per patient—identified via RF and SHAP analysis. For each subtissue cluster, we intersected the biomarker lists of all AD samples within that cluster to find genes that were consistently important across those individual samples. This process produced shared biomarker sets, varying in size from 4–80 genes per subtissue (Figure 3a). Such variation underscores the distinct pathobiology within different brain subtissues.

**Figure 3.**
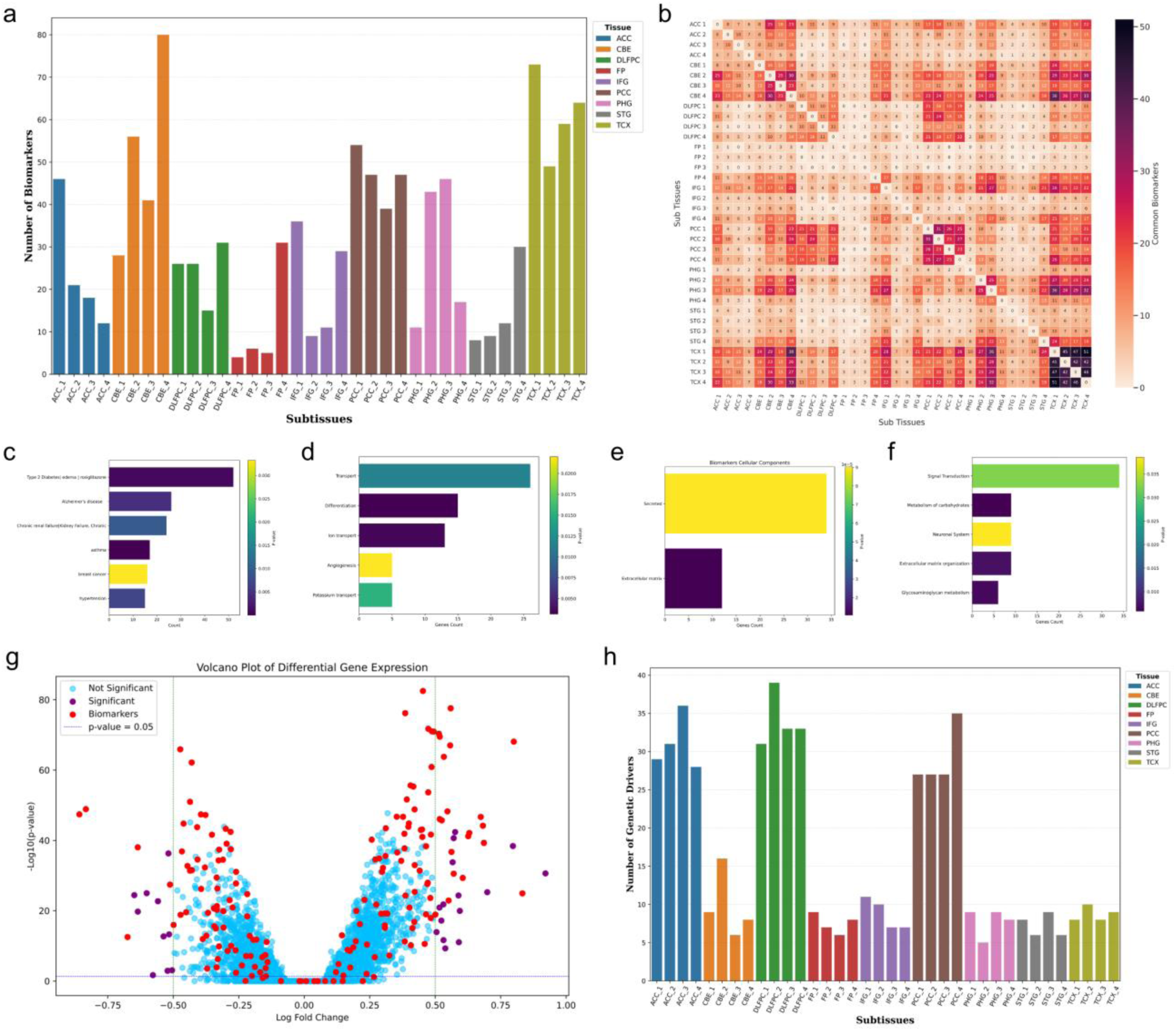
Subtissue-level biomarker discovery, functional enrichment, and genetic driver analysis in Alzheimer’s disease (AD). a) Bar plot of shared biomarkers (y-axis) identified by intersecting patient-level gene sets within each of the 36 subtissues (x-axis), ranging from 4–80 per subtissue/cluster. b) Heatmap showing the overlap of biomarker sets across different subtissues. Darker cells depict a greater degree of shared genes. c) Disease enrichment analysis (top six diseases) is based on the union of subtissue-specific biomarkers, highlighting comorbid or mechanistically related conditions (e.g., type 2 diabetes, chronic renal failure, and breast cancer). d–f) Representative Gene Ontology (GO) and Reactome enrichment results, demonstrating that the aggregated biomarkers are enriched in processes related to cell transport, differentiation, angiogenesis, extracellular matrix organization, and carbohydrate metabolism. g) Volcano plot from differential gene expression analysis comparing AD vs. control samples (|log2-fold change| > 0.5, and Bonferroni-adjusted *p* < 0.05). Biomarkers previously identified by the machine learning approach (colored points) show varied expression patterns, emphasizing the synergy between statistical and ML-driven methods. h) Analysis of 96 well-known AD-associated genes in genome-wide association study (GWAS) data, illustrating a bar plot of subtissue-specific genetic drivers. Statistical tests showed significant differences in mutation rates compared with background frequencies, underscoring the heterogeneous genetic architecture underlying AD pathophysiology across distinct brain tissues.

Combining the biomarker sets from all 36 subtissues produced 183 unique genes in total. A complete list of these genes, along with per-subtissue summary statistics, is provided in Supplementary Table 1. While Figure 2a confirms that overall transcriptomes segregate by tissue, the SHAP-derived biomarker sets isolate disease-relevant signals after controlling for baseline regional expression. This distinction allows us to pinpoint mechanisms uniquely perturbed in, e.g., the cerebellum versus the neocortex rather than merely reiterating constitutive tissue differences. A heatmap of these biomarkers highlights overlapping genes in pairwise subtissue comparisons (Figure 3b), revealing that some gene sets are highly tissue specific, whereas others span multiple subtissues.

#### 3.3 Functional Enrichment Confirms the Importance of Subtissue Biomarkers

To evaluate the biological significance of these 183 aggregated biomarkers, we performed GSEA. The full per-subtissue breakdown (e.g., the top genes in each subtissue) is available in Supplementary Table 1. This analysis showed associations of the gene set with diverse disease processes, including AD, type 2 diabetes, chronic renal failure, breast cancer, and hypertension (Figure 3c). These links may reflect shared pathways in metabolic disorders and neurodegeneration—such as insulin signaling deficits and dysregulated apoptosis implicated in AD (17,18)—or broader comorbid mechanisms. The significant enrichment of AD in the aggregated biomarker set shows that the RF–SHAP approach successfully prioritizes well-established AD genes, underscoring the reliability of the pipeline.

GO terms strongly enriched cell ion transport, differentiation, and angiogenesis, processes that are recognized as associated with the pathology of AD (16,19) (Figure 3d). The analysis also highlighted their involvement in cell secretion and extracellular matrix organization, processes critical in AD progression (15) (Figure 3e). Moreover, reactome pathway analysis revealed that biomarkers participate in signal transduction, carbohydrate metabolism, and extracellular matrix organization, further underscoring their comprehensive impact on cellular and molecular functions relevant to AD (15,20) (Figure 3f).

#### 3.4 Differential Expression Underscores Additional AD-Related Genes

As a complementary approach, we employed Limma (21) to compare AD vs. control expression levels across all tissues, identifying 2,347 differentially expressed genes (DEGs) with an absolute log2-fold changes > 0.5 at a Bonferroni-adjusted *p value* < 0.05 (Figure 3g). Among the 183 subtissue-specific biomarkers, 88 were significantly upregulated, and 72 were downregulated, whereas 23 did not meet the differential expression threshold yet still presented high SHAP importance. These 23 genes may exert regulatory effects through transcriptional or posttranscriptional mechanisms, as evidenced by several genes enriched in NOTCH1 signaling or reported to have high brain expression. Taken together, these findings highlight the added value of integrating ML-based feature selection with standard expression-level contrasts to capture a broader spectrum of AD-relevant genes.

#### 3.5 GWAS Data Pinpoint Subtissue-Specific Genetic Drivers

As AD progression is driven by both genetics and the environment, we next aimed to identify genetic signals in each subtissue. Using the individual-level genotypes available for all donors matched to our transcriptomic dataset, we first extracted the lead SNPs for 96 well-validated AD risk genes reported by (22). For every subtissue, we averaged each SNP’s risk-allele dosage and compared it with the AD cohort mean, showing clear subtissue-specific enrichment of AD variants and their parent genes (Figure 3h). Permutation testing confirmed that these dosage skews constitute distinct genetic signatures rather than random noise, highlighting regional variation in the disease’s genetic architecture. Hereafter, we refer to these genetic signatures (genes) as genetic drivers.

Integrating these subtissue-level genetic drivers with our previously derived transcriptomic biomarkers provides a more complete snapshot of AD pathology. However, phenotypic outcomes arise from interplay among multiple genes and pathways. To explore how genetic drivers might modulate the biomarker pathways, we subsequently constructed subtissue-specific gene‒gene interaction networks (see *Results section 4)*.

### 4) Network Analysis Prioritizes Critical Bottleneck Genes as Therapeutic Targets in AD

#### 4.1 Subtissue-Specific Gene‒Gene Interaction Networks Prioritize Key Mediators

To investigate how biomarkers and genetic drivers converge within each brain tissue, we constructed subtissue-specific gene‒gene interaction networks (details in *Methods section 5*). Each network was derived from RNA-seq profiles of AD samples in each subtissue and represents a coexpression network where edge weights correspond to Pearson correlation coefficients calculated between gene expression levels of samples. Only edges with correlation coefficients above a certain threshold were retained, ensuring that the network captures the strongest coexpression patterns among genes.

In these networks, biomarkers (orange nodes) and genetic drivers (green nodes) are typically not directly connected. Instead, they are linked through intermediate genes (blue nodes), referred to as "message-passing" genes, which mediate signaling cascades influencing AD progression and affecting the phenotype (Figure 4b). These message-passing genes were identified by computing the shortest paths between all pairs of subtissue-specific biomarkers and genetic drivers within each network. This approach selects genes that serve as critical connectors, potentially playing a key role in regulatory pathways, rather than simply identifying genes on the basis of high correlation or coexpression strength alone.

**Figure 4.**
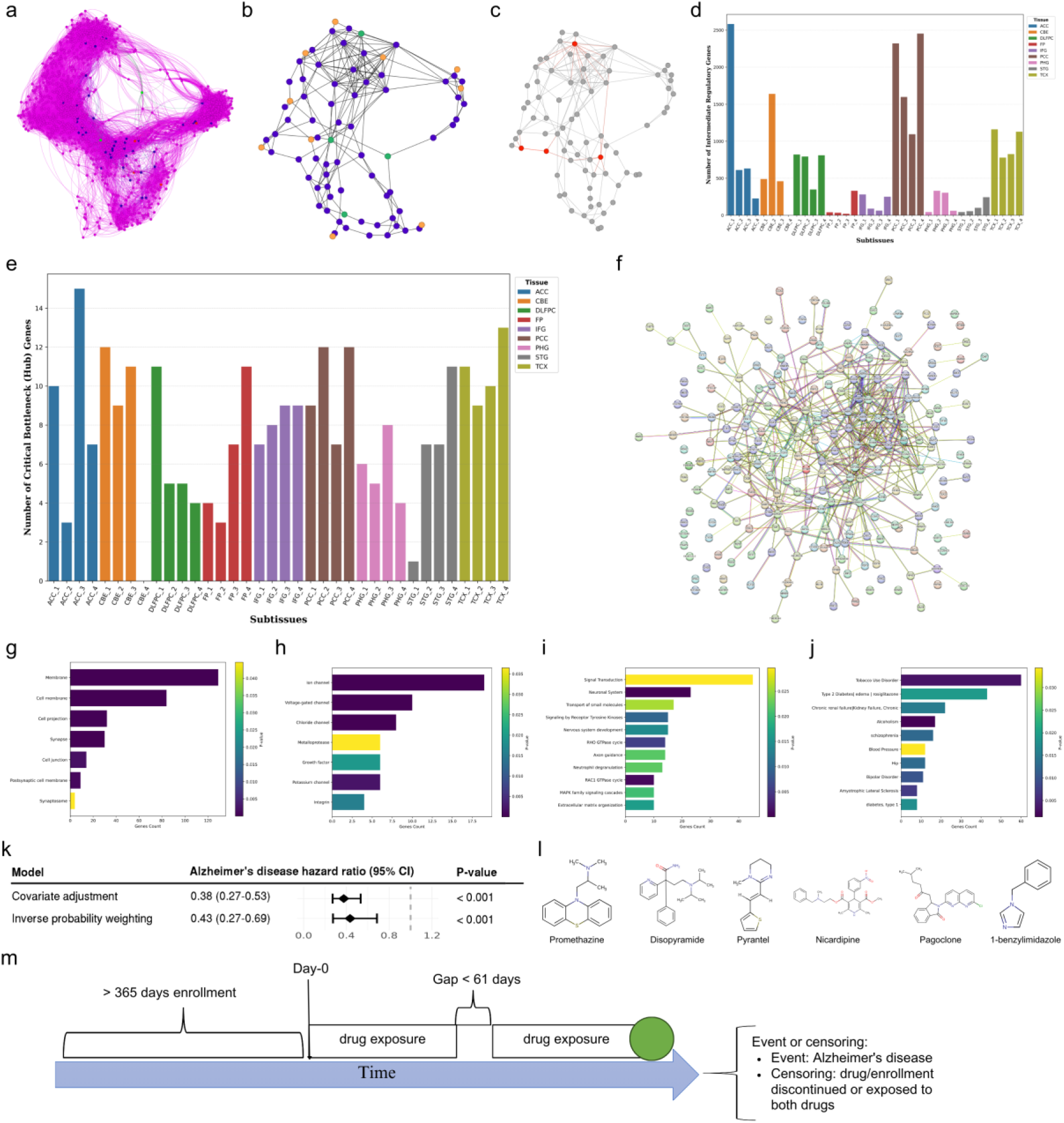
Subtissue-specific gene‒gene interaction networks, identification of critical bottleneck genes, and their functional relevance in Alzheimer’s disease. a) Weighted gene co-expression network (magenta) from a representative neocortex subtissue, capturing all genes expressed in that cluster. b) The same network filtered to include only biomarkers (orange nodes), genetic drivers (green nodes), and intermediate “message-passing” genes (blue nodes). c) Highlighting hub genes (red nodes) identified by multiple centrality measures (e.g., degree, betweenness, PageRank). d) Bar chart showing the number of intermediate message-passing genes for each of the 36 subtissues; one was excluded for insufficient samples. e) Variations in the counts of critical bottleneck genes (i.e., highly ranked novel hub genes) across subtissues, reflecting regional heterogeneity. f) STRING-based protein–protein interaction map of the prioritized 262 bottleneck genes, illustrating their interconnectivity in the broader human proteome. g–j) Functional enrichment analyses of these bottleneck genes, including cellular components (g), molecular functions (h), reactome pathways (i), and disease associations (j). The enriched terms highlighted roles in membrane localization, synapse organization, ion channel activity, signal transduction, cytoskeletal regulation, and various comorbid conditions such as type 2 diabetes. k) The forest plot displays hazard ratios (HRs) from two analytical approaches: covariate-adjusted Cox model (HR = 0.38) and the inverse probability of treatment weighting sensitivity analysis (HR = 0.43), both of which compare promethazine to cyproheptadine exposure and demonstrate consistent protective associations. l) Molecular structures of the six repurposed candidate drugs. m) Illustration of real-world pharmacoepidemiologic study design.

A representative example from one neocortex subtissue is shown in Figs 4a–4c. Figure 4a displays the entire subtissue coexpression network, whereas Figure 4b shows the isolates of only the biomarkers, genetic drivers, and their connecting intermediate message-passing genes. Figure 4c further highlights high-centrality (hub) genes in red, emphasizing their potential importance. Figure 4d illustrates the variability in the number of message-passing intermediate message-passing genes across different subtissues.

#### 4.2 Prioritizing Critical Bottleneck Genes through Centrality Metrics

In each filtered subtissue-specific gene‒gene interaction network, which included only biomarkers, genetic drivers, and intermediate message-passing genes, we ranked the constituent genes by multiple centrality metrics (e.g., betweenness, and closeness). These metrics assess a gene’s connectivity and influence within the network, pinpointing those with the most significant roles in mediating cellular communication within the subtissue network. Genes consistently scoring high in connectivity and low in clustering were designated candidate hub genes, as they appear pivotal in bridging communication between biomarkers and genetic drivers within the filtered network. This initial analysis yielded 22–30 candidate hub genes per subtissue.

To prioritize underexplored therapeutic possibilities, we then filtered out well-characterized AD genes (a gene set selected based on literature citation counts) from each hub gene list, focusing on less-studied but higher-ranked genes. These critical bottleneck genes may offer novel insights into AD pathophysiology and could serve as new therapeutic targets. The number of critical bottleneck genes varied across subtissues (Figure 4e), reflecting the heterogeneous genetic architecture of AD in different brain tissues. A complete listing of these bottleneck genes—including subtissue affiliations and annotation details—can be found in Supplementary Table 1.

Ultimately, unifying the critical bottleneck genes from all subtissues yielded 262 unique drug target candidates. Figure 4f illustrates how these bottlenecks interact in the broader human proteome (23). By integrating biomarkers, genetic drivers, and their bottleneck genes, this set underscores a systems-level perspective on AD pathology and suggests new directions for drug discovery and therapeutic interventions.

#### 4.3 Functional Enrichment Links Bottleneck Genes to Core AD Pathobiology

To evaluate the biological relevance of the 262 consolidated bottleneck genes, we performed GO and reactome enrichment analyses (Figs. 4g–4j). Most of these genes encode membrane or synaptic junction proteins involved in ion transport, adhesion, and differentiation—key processes linked to AD pathology (16,24). Other enriched functions included extracellular matrix organization and cellular secretion, both of which have been implicated in AD progression (15,25).

Reactome analysis highlighted processes such as signal transduction, nervous system development, small-molecule transport, actin cytoskeleton regulation, and axon guidance—all relevant to the molecular mechanisms of AD (16,25,26). Notably, cytoskeletal dysregulation has gained attention for its role in synaptic integrity (26), indicating that it is a potential area of further investigation.

Moreover, many of these bottleneck genes are associated with additional diseases—ranging from diabetes to neuropsychiatric disorders—suggesting shared mechanisms and underlining broader implications for comorbidities in AD. Together, these findings suggest that the identified bottleneck genes warrant deeper exploration as candidate targets for pharmacological intervention, which we address in the subsequent section on drug repurposing.

### 5) Drug Repurposing Suggests Multitargeted Strategies for AD Treatment

#### 5.1 Knowledge-Graph Analysis Discovers FDA-Approved Drugs Targeting Bottleneck Genes

To identify FDA-approved drugs capable of targeting multiple of the bottleneck genes (262), we utilized the Bioteque biomedical knowledge graph (10). This resource integrates over 150 data sources into a comprehensive network, containing more than 450000 biological nodes—such as 20,108 genes and 137396 compounds—and 30 million relationships, which include drug–gene, gene–disease, and other cross-domain interactions. Bioteque provides precomputed embeddings for the nodes, which capture the structure of the network, which includes both direct and multistep relationships (e.g., drug–gene–gene relationships). We systematically evaluated potential gene-drug interactions by calculating pairwise cosine similarities between the embeddings of our bottleneck genes and drugs included in the network. Drugs showing high similarity to multiple bottleneck genes were prioritized as promising multitarget therapeutic candidates for AD.

The Bioteque knowledge graph (10) connects drugs and genes through a network of biological relationships. We explored these relationships via "metapaths"—one-step pathways (drug-interacting-gene) and multistep pathways (e.g., compound-interacting-gene-coexpressing-gene). These multipaths reveal potential drug-gene relationships that go beyond direct interactions. We selected four specific metapaths, each representing a different way in which drugs and genes might be related biologically (see *Methods section 6* for details). For each metapath, we measured how similar each of the 262 bottleneck genes was to every compound in Bioteque via cosine similarity—a score that shows how closely related their numerical profiles (embeddings) are. We retained the top five compounds per gene on the basis of these scores.

Combining the results from all four metapaths yielded 2355 high-confidence drug-gene pairs, covering 225 bottleneck genes and 1,600 unique compounds. To narrow this down, we applied two filters: (1) a cosine similarity score of 0.65 or higher (see Supplementary Figure 1a), indicating a strong connection, and (2) drugs that target at least two bottleneck genes, suggesting a broader impact on AD (details in *Methods section 6*).

This process identified six FDA-approved drugs: pyrantel (5 targets), promethazine (3 targets), disopyramide (3 targets), nicardipine (2 targets), 1-benzylimidazole (2 targets), and pagoclone (2 targets). Figure 4l shows the molecular structures of these candidates. These drugs are promising for AD research because of their ability to affect multiple disease-related genes and pathways.

Three of these candidates (disopyramide, nicardipine, and 1-benzylimidazole) are predominantly cardiovascular agents that address conditions such as arrhythmias or hypertension. Pyrantel and promethazine exhibit anti-inflammatory or neuromodulatory properties, suggesting their relevance for AD interventions. Pagoclone, an anxiolytic partial gamma-aminobutyric acid (GABA)-A receptor agonist, presents a novel therapeutic avenue without the sedative drawbacks of related compounds. The following subsections describe each agent’s pharmacologic attributes and their potential impact on AD pathology.

#### 5.2 Predicted Repurposed Drugs Show Diverse but Similar Pharmacological Profiles

Our analyses highlighted six FDA-approved drugs (Figure 4l) —promethazine, disopyramide, pyrantel, nicardipine, pagoclone, and 1-benzylimidazole — that, despite disparate primary indications, converge on mechanisms potentially relevant to AD. Below, we summarize their documented pharmacological actions and AD-related rationales.

Promethazine (Phenergan) is a first-generation H1-antihistamine and an antagonist of the D2 dopamine receptor with strong sedative effects and moderate anticholinergic properties and is used for several allergic reactions (27). It also exhibits N-methyl-D-aspartate (NMDA) receptor antagonism, potentially conferring neuroprotective benefits by reducing excitotoxic damage. Clinically, promethazine is widely used for allergic symptoms, nausea, and adjunctive management of psychosis-induced aggression (28). *In vitro*, promethazine scavenges reactive oxygen species and modulates genes such as COX-2, suggesting anti-neuroinflammatory properties (29). A screen showed that promethazine can bind amyloid-β plaques in postmortem tissue, suggesting the possibility of modifying or imaging amyloid pathology (30). Additionally, when it is used alongside haloperidol for rapid tranquilization, it may also alleviate agitation or aggression in AD patients (28).

Disopyramide (Norpace), a class Ia antiarrhythmic agent, is employed to manage ventricular tachycardia and hypertrophic cardiomyopathy, reducing myocardial contractility and potentially benefiting neurological conditions through the modulation of voltage-gated sodium channels (31). While early AD is often accompanied by aberrant network hyperactivity or subclinical seizures, a sodium-channel blocker could stabilize firing rates and prevent excitotoxic damage (32).

Pyrantel, which is used to treat several worm infections, is a depolarizing neuromuscular blocker that mimics acetylcholine at nicotinic receptors in parasitic nematodes. It is FDA-approved for treating pinworm and hookworm infections (33). By activating nicotinic acetylcholine receptors, pyrantel can bolster synaptic plasticity and dampen neuroinflammation, aligning with cholinomimetic strategies in AD (34). Additionally, similar nicotinic receptor agonists (e.g., varenicline) have shown cognitive or anti-inflammatory benefits in models, reducing AD pathology (34).

Nicardipine (Cardene IV), an FDA-approved drug used to treat high blood pressure, is a dihydropyridine calcium-channel blocker that reduces Ca²⁺ influx into vascular smooth muscle, improves cerebral perfusion and lowers blood pressure (35). By preventing Ca²⁺ overload and increasing cerebral blood flow, nicardipine may protect neurons from ischemic and inflammatory damage (36). Epidemiological data suggest that dihydropyridines (nifedipine, nilvadipine) slow cognitive decline via antiamyloid or vasoprotective mechanisms (37). nicardipine is used for acute stroke management, implying potential synergy in AD, where chronic hypoperfusion accelerates pathology (38)

Pagoclone is a nonbenzodiazepine partial GABA-A receptor agonist designed for anxiety and tested for stuttering. Its partial agonism reduces anxiety with less sedation or dependence than full benzodiazepines do (39). Mild GABA enhancement can quell neuronal hyperexcitability, which is frequently observed in early AD, without the cognition-impairing effects of benzodiazepines (40). Moreover, the anxiolytic profile of pagoclone may help manage agitation or anxiety in AD patients (40).

1-Benzylimidazole, which is recognized for its positive inotropy and thromboxane A₂ synthase inhibition, has been explored as a cardiotonic and anti-inflammatory agent (41). Thromboxane A₂ is not only a platelet aggregator but also a proinflammatory mediator that can be produced in the brain. By inhibiting thromboxane A₂ production, 1-benzylimidazole may enhance cerebral perfusion and reduce proinflammatory eicosanoid levels, addressing microvascular deficits in AD patients (42).

Although each candidate poses distinct caveats—anticholinergic risks (disopyramide, promethazine) or uncertain central nervous system (CNS) penetration (pyrantel)—they share mechanisms potentially relevant to multiple AD pathways. Promethazine and pagoclone may mitigate neuropsychiatric symptoms and excitotoxicity, whereas nicardipine and 1-benzylimidazole target vascular and inflammatory processes, and pyrantel can bolster nicotinic cholinergic function. However, the potent antimuscarinic profile of disopyramide demands caution. Overall, these findings underscore the value of multitarget drug screening in AD and call for further research to determine whether such repurposed agents can meaningfully alter disease trajectories or symptom burden.

#### 5.3 Drug–Entity Network Suggests Convergent Pathways with Potential AD Relevance

To expand on each compound’s potential mechanisms, we extracted node embeddings from the Bioteque knowledge graph (10) and computed cosine similarities between each repurposed drug and the nodes of another graph entity, namely molecular functions. Retaining the top five hits per compound allowed us to pinpoint commonly repeated features that potentially underlie shared pharmacological effects.

Several compounds clustered around metabolic or contractile pathways—such as glycogen breakdown (glycogenolysis) and striated muscle contraction—have long been viewed as muscle-specific processes but are now recognized in AD because astrocyte-regulated glycogen metabolism affects neuronal signaling (43) and muscle-derived factors influencing hippocampal network development (44). We also identified repeated hits for processes such as DNA methylation and calcium-dependent regulation, which are known contributors to neuroinflammation and synaptic dysfunction in AD (45,46). Hence, the network-level findings suggest the multitarget potential of our candidate drugs. Recognizing these convergent pathways sets the stage for further validation: in the next section, we examine whether one of these candidates (promethazine) indeed demonstrates real-world benefits against the incidence of AD in large-scale real-world healthcare data.

### 6) Pharmacoepidemiologic Validation: Promethazine Shows a Reduced AD Risk Compared with an Active Comparator

To validate our drug repurposing predictions in real-world clinical settings, we conducted a retrospective pharmacoepidemiologic study using Optum’s de-identified Clinformatics® Data Mart (2007–2021). We employed an active comparator design to minimize confounding by indication—a critical consideration when studying drug effects in observational data. Rather than comparing promethazine users to nonusers (which could introduce bias due to different underlying health conditions), we compared promethazine exposure to cyproheptadine exposure, as both are first-generation H1-antihistamines with similar clinical indications but potentially different neurological effects (47).

After applying the inclusion criteria and wash-out periods (Figure 4m), our final cohort comprised 353856 individuals with promethazine exposure and 10877 with cyproheptadine exposure. This active comparator approach helps control for confounding factors related to allergy treatment needs while isolating the specific effects of promethazine on AD risk. The demographics of the study population are presented in Supplementary Table 2.

Using two complementary analytical approaches, both demonstrated considerable protective associations for promethazine compared with cyproheptadine. In the primary covariate-adjusted Cox proportional hazards model, promethazine exposure was associated with a 62% reduction in AD risk (hazard ratio (HR) = 0.38, 95% CI: 0.27-0.53, *p* < 0.001). A sensitivity analysis using the inverse probability of treatment weighting to further balance baseline characteristics between exposure groups yielded consistent results (HR = 0.43, *p* < 0.001) (Figure 4k). The consistency between these two analytical approaches—direct covariate adjustment and propensity score-based weighting—strengthens confidence in the observed protective association. The complete model coefficients and demographic characteristics are provided in Supplementary Tables S1-S3.

This considerable association between promethazine exposure and a reduced risk of AD, as evidenced by our robust covariate-adjusted Cox model, suggests potential neuroprotective properties that warrant further investigation. These findings could inform future clinical trials and healthcare strategies aimed at integrating promethazine into broader AD management protocols.

## Discussion

In this study, we introduce PRISM-ML, an integrated analysis pipeline that combines interpretable ML with multiomics data to explore the molecular complexity of AD. By analyzing transcriptomic and genomic profiles from the same patient cohort, we identified patient-level biomarkers and 36 subtissues across nine brain regions, highlighting the localized molecular diversity in AD.

We constructed subtissue-specific gene–gene interaction networks, identifying 262 high-centrality bottleneck genes as potential drivers of AD progression. Using a knowledge-graph approach, we predicted six FDA-approved drugs with multitarget potential, offering a promising strategy for therapeutics addressing multiple disease pathways. Our approach extends beyond previous studies by integrating multiple brain tissues, genomic variation, and real-world pharmacoepidemiologic validation.

Despite these advances, our study has limitations. It focused on transcriptomic and genomic data, excluding other omics layers like proteomics or metabolomics. One subtissue cluster lacked sufficient samples for reliable network construction. The use of bulk RNA-seq cannot resolve cell type-specific contributions, an area for future single-cell studies. Since samples in each tissue are contributed by a distinct cohort, formal batch-correction was not performed; any global adjustment would therefore downweight true regional differences—the very signal the study aims to characterize. Additionally, our drug repurposing relies on association-based knowledge graph embeddings, which require further validation to confirm interaction types (agonistic or antagonistic). However, our pharmacoepidemiologic findings on protective effects of promethazine highlight the potential of this approach.

Our research has several strengths that underscore its significance in AD research. First, by leveraging SHAP (8) values, we address the interpretability gap often associated with ML models, clearly illustrating each gene’s contribution to the classification of each AD sample. Second, our focus on subtissue-specific changes rather than broad tissue-level analyses enabled a more precise dissection of the regional heterogeneity of AD. Third, integrating multiomics data from the same cohorts offers a unified perspective on genetic and expression-level changes. Fourth, the LOOCV strategy ensures robust and personalized findings, addressing patient-specific disease manifestations. Finally, prioritizing novel bottleneck genes, rather than reiterating the well-studied AD genes, opens new therapeutic avenues.

Our GSEA revealed underappreciated pathways, such as synaptic function and actin cytoskeleton regulation, offering potential new targets. We also predicted six FDA-approved drugs (Figure 4l) with strong multitarget potential based on their interactions with critical bottleneck genes. While their primary uses differ, their shared mechanisms suggest relevance for AD.

Three of the six predicted drugs—disopyramide, nicardipine, and 1-benzylimidazole—are used primarily for cardiovascular conditions, whereas promethazine, pagoclone, and pyrantel modulate neuronal or neuromuscular pathways. Mechanistic overlaps suggest that promethazine and pagoclone may alleviate neuropsychiatric symptoms and excitotoxicity, whereas nicardipine and 1-benzylimidazole could address vascular and inflammatory dysregulation, and that pyrantel might bolster cholinergic function. Their side effects, particularly anticholinergic properties, require careful consideration.

The real-world evidence from our pharmacoepidemiologic study supports the protective effect of promethazine against AD. Using an active comparator design with cyproheptadine, another first-generation H1-antihistamine, our analysis of 364733 individuals demonstrated that promethazine exposure was associated with 62% risk of AD (HR = 0.38, 95% CI: 0.27-0.53, *p* < 0.001). This finding was further validated through sensitivity analysis via the inverse probability of treatment weighting (HR = 0.43, *p* < 0.001). These findings should be interpreted cautiously due to potential confounding and data limitations.

While these results are promising, they should be interpreted within the context of certain limitations, including potential unmeasured confounding effects, possible misclassification in health insurance records, and limited generalizability to populations outside commercial or Medicare Supplemental health insurance plans. Notably, healthcare claims data are vulnerable to misclassification of mild or early AD, which can bias hazard ratios. Despite these limitations, our active comparator design represents a methodological strength that distinguishes this study from simple exposure-versus-unexposed comparisons. By comparing two drugs with similar indications but different mechanistic profiles (47), we reduced the likelihood that the observed differences reflected underlying patient characteristics rather than drug-specific effects. The consistency of the results across the two analytical methods (direct covariate adjustment and inverse probability of treatment weighting) further supports the robustness of our findings.

One study (27) has shown that a high anticholinergic load can worsen confusion, causing delirium in older adults; therefore, chronic use of promethazine in AD patients demands careful risk–benefit assessment. Future studies should validate these results in more diverse cohorts and incorporate rigorous causal-inference methods to address residual confounding.

Moving forward, our findings set the stage for deeper investigation into these identified candidates and their biological interactions. Rigorous *in vitro* and *in vivo* assays can clarify whether each agent’s mechanism—agonistic or antagonistic—truly counter AD pathologies. In parallel, novel computational methods, including neural networks and transformer-based models with interpretability tools (49), could refine biomarker discovery, whereas expanded omics data layers (e.g., proteomics, metabolomics, and epigenetics) might yield a fuller picture of the molecular architecture of AD. Future investigations should incorporate single-cell and spatial transcriptomics, which can resolve the cell type-specific disruptions masked by bulk RNA profiling, leading to refinement of our understanding of molecular architecture of AD. Ultimately, PRISM-ML bridges foundational research and clinical application, paving the way for multipathway therapeutics tailored to individual profiles.

## Conclusion

In this work, we developed PRISM-ML, an interpretable ML analysis pipeline integrated with multiomics data (bulk RNA-seq and genomics) to illuminate key molecular mechanisms underlying AD. By identifying patient-level biomarkers and constructing subtissue-specific gene‒ gene interaction networks, we prioritized critical bottleneck genes and highlighted new paths for therapeutic intervention. Our data-driven knowledge graph-based approach led to the discovery of six repurposable drugs that target multiple AD-relevant pathways simultaneously, marking a shift from conventional single-target strategies. Real-world evidence, including pharmacoepidemiologic validation of promethazine, underscores the clinical promise of our findings, although further experimental and clinical studies are needed to refine the mechanistic details and therapeutic efficacy.

As precision medicine gains momentum, our integrated analysis pipeline—combining molecular signatures, knowledge graph-based drug repurposing, and pharmacoepidemiologic validation— offers a powerful template for unraveling the heterogeneity of complex diseases and accelerating the development of targeted, multipathway drug treatments. While further experimental work is necessary to validate our findings, this study paves the way for transformative advancements in AD treatment and beyond, potentially accelerating the shift toward more personalized medicine in clinical practice.

## Material and Methods

### 1) Data Collection and Preprocessing

We integrated bulk RNA-sequencing and GWAS data, from three independent large-scale AD brain tissue studies: the Religious Orders Study and Rush Memory and Aging Project (ROSMAP) (11), the Mount Sinai Brain Bank study (MSBB) (12), and the Mayo RNA-seq study (MAYO) (13), which were obtained via the Accelerating Medicines Partnership - AD (AMP-AD) Knowledge Portal (https://www.synapse.org/Synapse:syn21241740 and https://www.synapse.org/Synapse:syn22264775) (50). In total, our curated dataset comprised 2105 postmortem brain tissue samples (1363 AD and 742 control) from 835 unique individuals (528 females and 307 males) collected from nine distinct brain tissues: anterior cingulate cortex (ACC), CBE, dorsolateral prefrontal cortex (DLFPC), FP, inferior frontal gyrus (IFG), PCC, parahippocampal gyrus (PHG), superior temporal gyrus (STG), and TCX. Table 1 summarizes the distribution of samples and subjects across the nine brain tissues.

As our objective was to discover subtissue-specific biomarkers and genetic drivers, we treated tissue—not cohort—as the primary biological variable. In the AMP-AD collection, every brain tissue is contributed by only one cohort (ROSMAP, MSBB, or Mayo), making the “cohort” and “tissue” inseparable. Therefore, without performing global batch effect correction, we retained the log-scaled, within-tissue–normalized expression data from each cohort, which were also regressed out for nonbiological covariates (e.g., batch number, library size, RNA integrity, postmortem interval, age, sex) while leaving the diagnosis variable unchanged.

We utilized individual-level genotype data from the same donors whose brain tissue provided the RNA-seq data, ensuring matched genomic and transcriptomic profiles from identical individuals. Genotypes were coded as dosages (0–2). The AD-associated genetic variants analyzed were selected based on prior GWAS meta-analyses, specifically focusing on a study (22) that identified 96 genes significantly enriched for AD association through large-scale GWAS summary statistics.

In this study, each postmortem RNA-seq + GWAS profile comes from a unique donor; therefore every sample represents an independent patient, and we use ‘patient-specific’ (synonymous with “per-donor” or “sample-level”) profiles throughout the paper.

### 2) ML Classification Model and Model Interpretation

To identify patient-specific (per-donor) biomarkers that distinguish AD samples from control samples, we trained RF classifiers implemented in Python (v3.10.6) via scikit-learn (v1.4.0) (51). The top 5000 highly variable genes across all samples served as input features and were selected after comparative testing of alternative feature sets (3000–12000 genes). This feature size was chosen because it maximized model performance and yielded significant enrichment of AD-related genes in subsequent GSEA (see *Methods section 3*). Notably, established AD-associated genes (e.g., APOE) were omitted automatically owing to minimal expression variation across samples.

To ensure robust performance, we optimized the RF binary classifier (AD samples vs control samples) via a grid search with 10-fold stratified cross-validation. This process systematically evaluated combinations of hyperparameters, yielding the optimal configuration: {n_estimators=722, max_depth=38, min_samples_split=5, min_samples_leaf=4, max_features=0.11}. With these parameters, the model achieved strong performance during cross-validation: an F1 score of 0.88, precision of 0.84, recall of 0.93, and receiver operating characteristic area under the curve (ROCAUC) of 0.904.

For the final evaluation, we employed a LOOCV strategy. In LOOCV, each sample in the dataset is withheld once as a test set, whereas the remaining samples are used to train the model. This ensures that the model’s performance is assessed on unseen data for every sample. For our dataset of 2105 postmortem brain samples, we trained one RF model per sample—totaling 2105 models— where each model was trained on 2104 samples and tested on a single withheld sample. The class-specific performance metrics of these RF models are presented in Table 2 below. The overall accuracy of the models was 0.88, reflecting robust predictive performance across the entire dataset.

To understand the gene-level contributions to the model’s predictions, we utilized TreeExplainer from the SHAP method (52). SHAP offers a unified measure of feature importance, quantifying the impact of each gene/feature on the classification decision of individual models and samples. Unlike the built-in feature importance of RF (mean-decrease-in-impurity), which only shows magnitude, SHAP provides unbiased, sample-specific explanations with both magnitude and direction. A positive SHAP value indicates that the expression of the gene pushes the model toward predicting AD, whereas a negative value pushes it toward the control class. This directional insight makes SHAP a better choice for interpreting the model’s decisions.

For every AD sample that the model classified correctly, we computed SHAP scores for the full set of 5000 input genes. Plotting the cohort-wide distribution of SHAP values revealed a clear elbow at SHAP = 0.00026 (Supplementary Figure 1b): above this point the curve flattens, indicating a small fraction of genes that together explain the bulk of the model’s decision. We therefore label genes with SHAP ≥ 0.00026 in a given sample as high-impact genes. This data-driven threshold produces a median of 175 genes per donor (interquartile range 152–222), and we refer to these sample-level lists as patient-specific biomarker sets.

In addition to identifying biomarkers, we utilized the *predict_proba()* function of our models to generate a classification predictive (probability) score for each sample. This score, ranging from 0 to 1, indicates the model’s confidence in classifying a sample, with higher values representing a greater likelihood of AD. We selected AD samples that were correctly classified with a probability exceeding the average probability of the AD class for subsequent clustering and analyses. This filtering step enhanced robustness without biasing downstream analyses.

### 3) Subtissue Identification and Subtissue-Specific Biomarkers and Genetic Drivers

#### 3.1 K-Means Clustering

To identify finer-scale molecular variations within each of the nine brain tissues, we applied K-means clustering to group AD samples on the basis of their gene expression profiles. For each tissue, we selected AD samples that were confidently classified by our RF model—specifically, those with predictive scores exceeding the average for the AD class (0.82). We then constructed a feature matrix for each tissue using these confidently classified AD samples. The features in this matrix were the union of all patient-level biomarker sets identified for the samples within that tissue, ensuring that the most relevant genes for distinguishing AD pathology in that region were included.

Using this tissue-specific feature matrix, we performed unsupervised K-means clustering with (k=4) to divide the samples into four distinct "subtissues." The optimal number of clusters was determined by evaluating silhouette scores, which measure how well separated and cohesive the clusters are, and the Calinski–Harabasz index (variance ratio criterion), which assesses the ratio of between-cluster to within-cluster variance. Higher values for these metrics indicate better-defined clusters. We tested k values from (2–10) and observed that k = 4 consistently produced high values for at least one of these metrics in every tissue (Supplementary Figure 2). Accordingly, we set k = 4 for all the tissues, yielding 36 subtissues across the nine brain tissues. Supplementary Table 1 summarizes the number of samples in each subtissue.

#### 3.2 Subtissue-specific Biomarker Definition

We intersected the patient-level biomarker lists of all AD samples within each of these subtissue clusters to define ‘subtissue-specific biomarker sets’ (ranging from 4–80 genes per cluster). Details of these gene sets and their associated statistics are available in Supplementary Table 1. Specifically, (1) each cluster (subtissue) contained a subset of AD samples, each with its own ∼175 patient-level biomarkers, and (2) we took the intersection (common genes) across these AD samples. Genes appearing in all AD samples of that subtissue were designated “subtissue-specific biomarkers.” This approach ensures that the resulting genes are consistently highly impacted in the same cluster of AD samples, highlighting regionally shared dysregulations. Subtissue-specific biomarker set sizes ranged from 4–80 genes, reflecting heterogeneity among subtissues. Notably, repeating the SHAP analysis on a diagnosis-shuffled label set produced no consistent subtissue-specific biomarker signatures.

#### 3.3 Identifying Subtissue-Specific Genetic Drivers

We curated a list of 96 well-characterized AD-associated genes (and their lead single nucleotide polymorphisms (SNPs)) from recent GWAS meta-analyses (Andrade-Guerrero et al., 2023). For each sample in our study, per-allele genotype data for these SNPs were extracted from the matched GWAS files.

To pinpoint tissue-specific genetic drivers, we computed mutation rate (frequency) of each SNP and allele frequency within every subtissue (AD samples only) and then performed the following: (1) we compared single nucleotide polymorphism (SNP) rates (e.g., minor allele frequency) against the full AD cohort as a background; (2) we performed Fisher’s exact tests, followed by empirical permutations, assessing whether a given enrichment of a SNP in a subtissue was greater than expected by chance; and (3) we assigned SNP-associated genes to a subtissue if their SNPs exhibited significant enrichment (*p* < 0.05 after multiple-testing correction). These genes, together with the subtissue-specific biomarkers, were subsequently used to construct gene‒gene interaction networks.

### 4) Statistical Analysis and Gene Set Enrichment

Differential gene expression was assessed via the Limma package (v3.62) in R (21). Genes with an absolute log2 fold change > 0.5 and a Bonferroni-adjusted *p* value < 0.05 were considered significantly different between the AD samples and the control samples.

Gene set enrichment analysis (GSEA) was performed via the DAVID database (14) to evaluate the biological functions of tissue- and subtissue-specific biomarkers. Enriched GO terms and Reactome pathways highlighted processes such as membrane binding, extracellular matrix organization, and ion transport—pathways that are critically involved in AD pathogenesis.

### 5) Gene-Gene Interaction Network Construction and Critical Bottleneck Genes

We employed WGCNA (7) in R to construct subtissue-specific gene‒gene interaction networks. For every subtissue, we chose a WGCNA soft threshold (soft-power 𝛽) that maximized both the mean connectivity and scale-free topology fit (R²≥0.85) via an adjacency matrix. Adjacency matrices were computed based on Pearson correlations from the gene expression profiles in each subtissue. To preserve biologically realistic network sizes, we retained the strongest edges (edge weights set based on Pearson correlation) until the final subtissue network had an edge density of ∼0.5%, aligning with typical protein–protein interaction densities in the human proteome (23). This procedure yielded networks that are comparable in their topology (scale-free degree distribution) without enforcing an identical number of edges.

We chose not to normalize edge density across the 36 subtissue networks for three methodological reasons: (1) scale-free baseline: each subtissue’s network was already scale-free due to individually selected soft thresholds; (2) intranetwork ranking: bottleneck genes were ranked within each subtissue network using flow centrality, making cross-network normalization unnecessary; and (3) biological variation: differences in network density reflect genuine biological differences between subtissues (e.g., cellular composition and transcriptional coordination), which we aimed to preserve rather than suppress. Notably, coexpression does not imply directionality or causality, particularly in bulk RNA-seq data, where cell type composition and shared variance can introduce confounding factors.

For each subtissue (as defined by K-means clustering), we then compiled two gene sets and flagged them in the network: (1) Subtissue-specific biomarkers—genes consistently flagged by SHAP across all AD samples in that cluster—and (2) genetic drivers—the subset of 96 AD-associated GWAS genes that showed significant enrichment in the same subtissue. The combined gene set ensured that disease-relevant signatures (biomarkers/drivers) were flagged. All networks were visualized with the String database (23) and Gephi (53).

To probe how these two classes interact, we computed the shortest paths connecting every biomarker to every driver within each subtissue network. Every node lying on at least one such path was tagged as a potential “message-passing” intermediate, illuminating genes that may mediate communication between primary biomarkers and genetic drivers. Figure 4b shows the number of intermediate ‘message-passing’ genes identified in each subtissue.

We then computed multiple centrality measures (e.g., betweenness centrality, PageRank, closeness, and eigenvector centrality) to quantify each gene’s importance in relaying network information. Genes consistently scoring in the top ranks across multiple metrics were designated as high-centrality “hub” genes, reflecting their importance in bridging multiple AD-relevant pathways. This step yielded 22–30 candidate hub genes per subtissue. We then focused on the hub genes ranked higher than the well-characterized AD genes (e.g., APOE) based on the literature citations, to explore novel candidates, yielding 1–15 critical “bottleneck” genes per subtissue (Figure 4e). Combining bottleneck gene lists across all 36 subtissues produced a nonredundant set of 262 candidate drug targets.

### 6) Drug Target Identification and Repurposing Drugs

Drug repurposing analysis was conducted via the Bioteque knowledge graph (10) which integrates 30 million relationships across 12 biomedical entity types. We systematically queried four metapaths to model compound-gene interactions: (1) CPD-int-GEN: direct pharmacological interactions (e.g., binding, inhibition), (2) CPD-int-GEN-cdp-GEN: compound effects mediated by codependent genes, (3) CPD-int-GEN-cex-GEN: compound-gene associations via coexpressed intermediaries, (4) CPD-int-GEN-ppi-GEN: compound effects propagated through protein interactors.

We focused solely on 262 bottleneck genes identified from our subtissue-specific interaction networks (see *Results*). For each metapath, node embeddings (128-dimensional vectors) were extracted, and cosine similarities between all 262 bottleneck genes and 137396 compounds were computed. We then retained only the top five compounds per gene (i.e., those with the highest similarity), thereby discarding low-confidence pairs and reducing noise.

We concatenated the reduced results from each route, resulting in a final dataframe of 2355 gene– drug pairs covering 225 bottleneck genes and 1600 unique drug entities. We then plotted the distribution of cosine similarities across these embedding pairs and selected the threshold of 0.65 where an elbow was observed (Supplementary Figure 1a). Next, we filtered out drugs that strongly interact with only a single bottleneck gene, as our aim was to highlight multitarget therapeutic leads. From this process, six Food and Drug Administration (FDA)-approved drugs were found to surpass both criteria—disopyramide, nicardipine, 1-benzylimidazole, pagoclone, promethazine, and pyrantel.

### 7) Pharmacoepidemiologic study

We used Optum’s de-identified Clinformatics Data Mart Database (Clinformatics, years: 2007-2021), licensed to Indiana University. Clinformatics is derived from a database of administrative health claims for members of large commercial and Medicare Advantage health plans. Informed consent was waived by the Indiana University Institutional Review Board.

We used an active comparator design by selecting cyproheptadine (10877 individuals) as the comparator for promethazine (353856 individuals), as both drugs are tricyclics and first-generation H1-antihistamines (47). We defined the initiation date (e.g., day 0) as the first date of exposure. We included individuals who were started on promethazine and/or cyproheptadine, were aged ≥60 years and had an enrollment period of ≥2 years. We excluded individuals who had both drug exposure on day-0, only 1-day of exposure, and missing gender information; as well as individuals with the following conditions prior to day 0: <365 days of enrollment, AD, dementia, AIDS, solid tumors without metastasis, lymphoma, and/or metastatic cancer. Figure 4m illustrates the pharmacoepidemiologic study design and cohort construction.

We used International Classification of Diseases (ICD) codes (e.g., ICD-9 3310, ICD-10 F00*, and ICD-10 G30*) to define AD (54,55). We defined the outcome as the time from day 0 to the first AD diagnosis date. We defined the censoring date as the earliest of discontinuation of exposure (e.g., the last date of exposure without exposure in the next 60 days), concurrent exposure to both drugs, or discontinuation from enrollment.

We collected covariates on demographics, including age, gender, race and index year of day-0. The demographics of the study population are provided in Supplementary Table 1. Additionally, we used the R package Comorbidity to define alcohol use disorder, blood loss anemia, cardiac arrhythmias, congestive heart failure, coagulopathy, chronic pulmonary disease, deficiency anemia, depression, diabetes status (uncomplicated and complicated), drug abuse, fluid and electrolyte disorders, hypertension status (uncomplicated and complicated), hypothyroidism, liver disease, obesity, other neurological disorders, paralysis, pulmonary circulation disorders, psychoses, peptic ulcer disease excluding bleeding, pulmonary circulation disorders, peripheral vascular disorders, renal failure, rheumatoid arthritis or collaged vascular disease, valvular disease, and weight loss. Moreover, we identified fall, hearing impairment/loss and vision impairment according to (56).

We used Cox models to estimate hazard ratios (HRs), 95% confidence intervals (CIs), and *p* values. First, we fitted a covariate-adjusted Cox model by adjusting all covariates. Second, we conducted a sensitivity analysis utilizing the inverse probability of treatment weighting with the propensity score. We used a logistic regression model to estimate the propensity score by controlling for all covariates. The Cox model included the exposure status of promethazine/cyproheptadine and used inverse probabilities of treatment as weights. The covariate-adjusted Cox model results are presented in Supplementary Table S3. Figure 4k shows the adjusted hazard ratios. All pharmacoepidemiologic analyses were conducted in R.

### 8) Statistics and Reproducibility

All analyses were performed with appropriate sample sizes and clearly stated statistical methods. We compiled 2105 postmortem brain tissue samples (1,363 AD cases and 742 controls) across nine brain tissues; one subtissue cluster was excluded from the gene‒gene interaction network construction because of the limited sample size (n=2). For predictive modeling, we employed an RF classifier (which does not require normally distributed data) and used a LOOCV analysis pipeline (two-tailed), reporting precision, recall, and F1 scores for AD sample and control sample classifications. A total of 5,000 highly variable genes served as the input feature set. We identified DEGs using Limma (v3.62) in R (two-tailed) with an absolute log2-fold change >0.5 and a Bonferroni-adjusted p value<0.05, thereby controlling for Type I error inflation. K-means clustering (k=4 per tissue) was validated via silhouette scores and Calinski–Harabasz indices (Supplementary Figure 2). Gene expression distributions were deemed suitably handled by the empirical Bayes approach of Limma; no additional normality checks were conducted given the large sample sizes per tissue (Table 1).

Subtissue-level genomic variation was examined via two-sided permutation tests of continuous SNP dosage data (n=96 AD-associated genes), comparing the mean allele dosage in a given cluster against the remaining samples (5000 random resamplings, *p*<0.05). We also performed two-tailed Fisher’s exact tests to detect enrichment of minor allele frequencies (categorical variables present in two datasets) for GWAS-derived variants, as chi-square tests were not appropriate for small subtissue samples. GSEA was carried out in DAVID (14) via hypergeometric tests with Benjamini correction (q value<0.05). Finally, our pharmacoepidemiologic analysis employed two-tailed Cox proportional hazards models (both covariate-adjusted and inverse probability–weighted) at α=0.05, which compared the incidence of AD among individuals exposed to promethazine and a matched active comparator; effect sizes are reported as hazard ratios with 95% confidence intervals and corresponding p values.

## Data Availability

The analyzed bulk RNA sequencing and genomics (GWAS) datasets of ROSMAP, MSBB, and MAYO cohorts can be accessed with consent via the AD Knowledge Portal (https://adknowledgeportal.org; accession No. syn21241740 and No. syn22264775) (50).

Optum’s de-identified Clinformatics® Data Mart Database is not publicly available (accessibility can be obtained from Optum: https://www.optum.com/en/). The complete underlying code for this study is available in PRISM-ML_AD GitHub repository and can be accessed via this link https://github.com/mmottaqii/PRISM-ML_AD.

## Funding

This project has been funded by the National Institute of General Medical Sciences of the National Institute of Health (R01GM122845), the National Institute on Aging of the National Institute of Health (R01AG057555, R21AG083302), and the National Science Foundation (2226183). The funders played no role in the study design, data collection, analysis and interpretation of the data, or the writing of this manuscript.

## Author contributions statement

M.M. prepared the data, implemented the algorithms, performed the experiments, analyzed the data, and wrote the manuscript; P.Z. performed healthcare data mining, and wrote the manuscript; L. X. conceived and planned the experiments, and wrote the manuscript.

## Ethics approval and consent to participate

The use of electronic health record (EHR) data for the pharmacoepidemiologic analysis was reviewed by the Indiana University Institutional Review Board (IRB), which designated the study as exempt from further review (no human subjects research). The dataset—Optum’s de-identified Clinformatics Data Mart Database—contains fully de-identified administrative claims for large commercial and Medicare Advantage health plan members, and informed consent was not required because all patient identifiers were removed prior to data access. Permission to use and analyze the de-identified dataset was obtained from the data provider. No additional consent to participate was needed for this secondary analysis of existing de-identified data. This study complies with ethical standards for retrospective analysis of de-identified data as per the Declaration of Helsinki and relevant regulations.

## Competing Interests Statement

All authors declare no financial or non-financial competing interests.

## Acknowledgements

The authors acknowledge the important contributions of three publicly available datasets, including the Religious Orders Study/Memory and Aging Project (ROSMAP), the Mount Sinai Brain Bank (MSBB), and the MAYO. We thank the participants of the ROS, MAP, MSBB, and Mayo for their time and participation.

## Notes

### Competing Interest Statement

The authors have declared no competing interest.

### Summary of Updates

Supplementary files were updated; additional technical details were added.

