## Supplemental material for "Integrating explainable AI with multiomics systems biology and EHR data mining for personalized drug repurposing in Alzheimer’s disease"

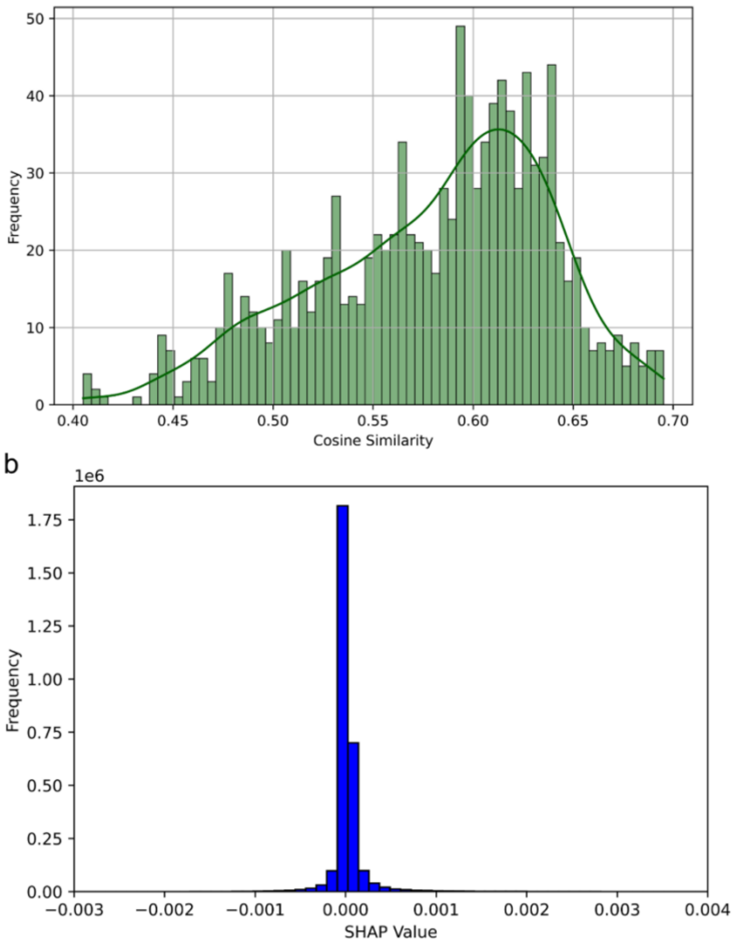


**Supplementary Figure 1. Data-driven cutoffs are used to retain high-confidence drug–gene links and high-impact genes (unique sample-level biomarkers identified by SHAP as a feature importance measurement method).**

a) Histogram of cosine-similarity scores for the 2355 drug–gene embedding pairs (128 dimensions) obtained by merging the four metapath searches (green kernel-density overlay). The distribution shows an "elbow" at a similarity of 0.65, which we adopt as the minimum score for a reliable drug–gene association.

b) Cohort-wide distribution of SHapley Additive exPlanations (SHAP) values computed for all 5,000 input genes in every correctly classified AD sample. The values cluster tightly around zero with a sharp fall-off above $2.6 \times 10^{-4}$, indicating a natural break that we use to define "high-impact" genes in each AD sample. Together, these thresholds focus subsequent analyses on the most biologically and pharmacologically meaningful signals.

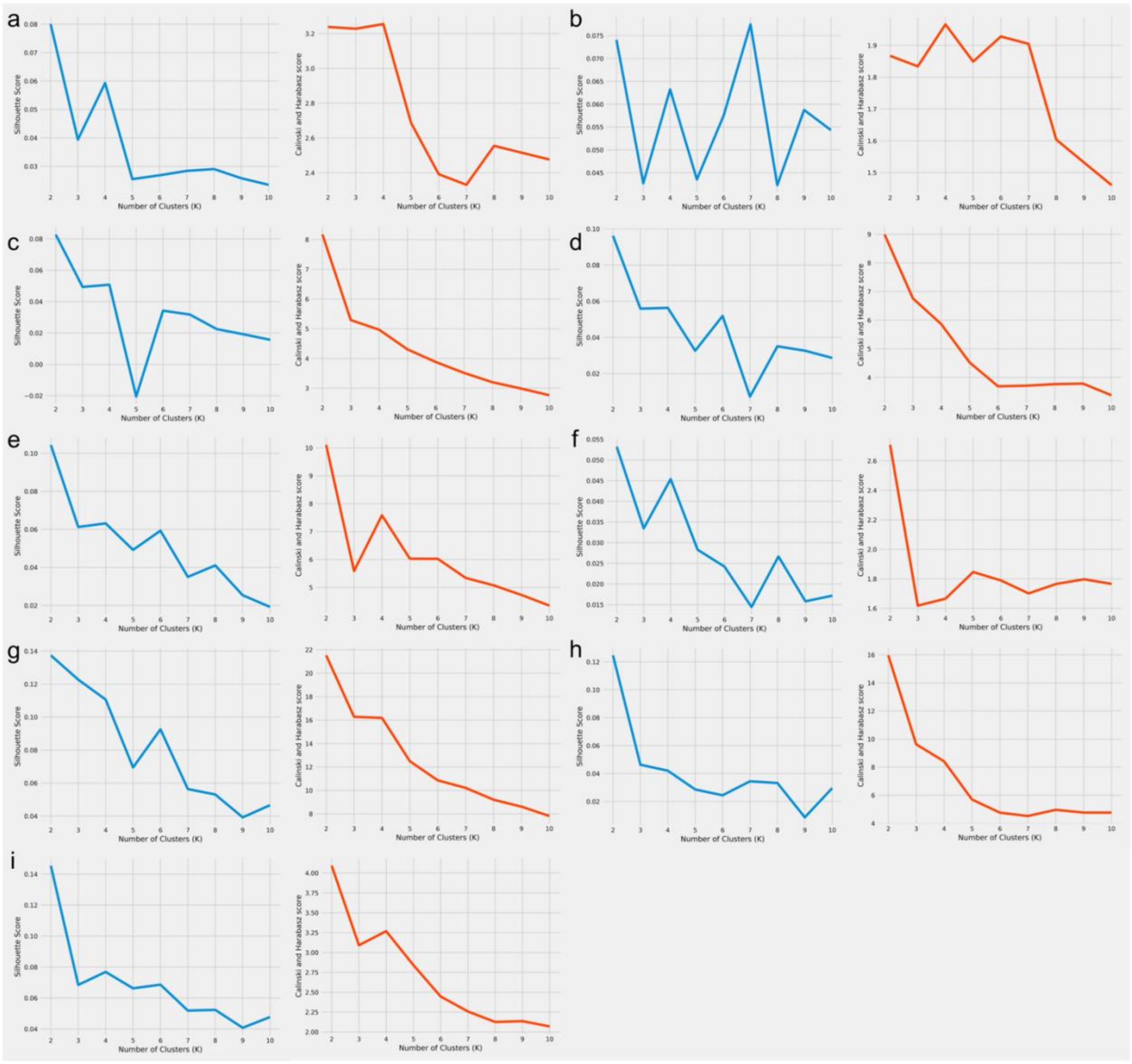

**Supplementary Figure 2. Selecting the optimal number of K-means clusters for each brain tissue.**

For each of the nine brain tissues analyzed (panels a–i), we varied the number of K-means clusters (k = 2–10) and quantified clustering quality with two complementary, scale-independent metrics. The silhouette score (blue line, left-hand plot in every panel) measures the cohesion and separation of individual samples (higher values denote better-defined clusters). The Calinski–Harabasz (C–H) index (orange line, right-hand plot) is the ratio between–cluster dispersion and within-cluster dispersion (higher values indicate tighter, better-separated clusters). In every tissue, k = 4 either maximized or lay on a clear local peak (“elbow”) for at least one of the two metrics, whereas larger k values showed diminished or unstable scores.
